# Leaf traits predict performance under varying levels of drought stress in cultivated sunflower (*Helianthus annuus* L.)

**DOI:** 10.1101/2023.03.06.531401

**Authors:** Ashley M. Earley, Kristen M. Nolting, John M. Burke

## Abstract

Drought is a major agricultural challenge and is expected to worsen with climate change. Exploring plant traits and how they respond to drought has the potential to improve understanding of drought tolerance and inform breeding efforts to develop more drought tolerant plants. Given their importance in plant-water relations, we explored variation and plasticity in leaf traits in response to water limitation in cultivated sunflower (*Helianthus annuus* L.). A set of four sunflower genotypes was grown under four different levels of water availability and leaf vein and stomatal traits were measured along with total biomass (as an indicator of performance), leaf mass per area (LMA), chlorophyll content, and various mass fraction traits related to resource allocation (e.g., leaf, root, and stem mass fraction). Traits exhibited numerous bivariate correlations within treatments that generally followed expectations based on the literature. For example, stomatal size and density were negatively correlated while stomatal density and vein length per area (VLA) were positively correlated. Most traits exhibited substantial plasticity, as evidenced by significant shifts in trait values across environments and multivariate analyses revealed differentiation in trait space across treatment levels. This included an overall reduction in growth/productivity in response to stress, accompanied by a shift in traits relating to gas exchange and hydraulics including stomatal and vein density (increased), stomatal size (decreased), and theoretical gsmax (increased). We found that variation in performance across treatments (estimated as total biomass) can be largely explained by a small number of putatively size-independent traits (i.e., VLA, stomatal length and density and LMA; *R*^2^ = 0.74). Moreover, on average, more extreme changes in VLA were associated with more extreme decreases in performance across environments. A small number of leaf traits can predict plant performance, with plasticity in VLA being the best predictor of changes in productivity.

## INTRODUCTION

Drought is a major agricultural challenge that limits plant growth and productivity worldwide (Munns, 2011). As climate change worsens, droughts are expected to increase in frequency and severity (Foley *et al*., 2011; Munns, 2011; IPCC, 2014). Moreover, the human population is expected to increase from a current size of 7.6 billion to 9.8 billion by 2050 (United Nations, 2017). This poses a challenge as the growing population size will increase food demand while much of our agricultural land will be increasingly impacted by environmental stresses including drought. To address this challenge, we need crop plants that are better able to withstand environmental challenges such as drought. A better understanding of the response of key traits to stress will facilitate efforts to develop more resilient crops. For example, given their influence on hydraulics and gas exchange, leaf anatomical traits such as stomatal and vein-related traits have been the subject of considerable interest in the context of drought (e.g., Bresta *et al*., 2018; Du *et al*., 2021; Lei *et al*., 2018). Interestingly, associations between these sorts of traits and relevant aspects of the environment have been observed both across and within species suggesting a possible role for such traits in adaptation to drier environments (e.g., Bresta *et al*, 2018; Lei *et al* 2018). For instance, in barley, plants with smaller and denser stomata as well as denser veins showed less dramatic changes in these traits under drought (Bresta *et al*, 2018).

When looking across species, vein density and stomatal density are positively correlated with each other (Sack and Scoffoni, 2013), while stomatal density and size are negatively correlated (Shahinnia *et al*., 2016; Doheny-Adams *et al*., 2012). This relationship between stomatal size/density and vein density (estimated as vein length per area; VLA) is important for optimization of water transport and transpiration (Fiorin *et al*., 2016; Bertolino *et al*., 2019). Consistent with a role for such traits in adaptation to dry environments, populations of plants occupying increasingly arid habitats exhibit a trend toward smaller and more dense stomata (Xie, Wang, and Li 2022; Dunlap and Stettler 2001). For example, across 19 *Protea repens* populations, stomatal density in a common garden was found to correlate negatively with summer rainfall levels. In addition to inherent differences between populations, these traits have also been shown to exhibit plastic responses to water limitation. For example, plants often shift toward increased VLA, higher stomatal density, and smaller stomata under drought vs. well-watered conditions (Sack and Scoffoni, 2013; Sun *et al*., 2014). These changes are thought to improve gas exchange by decreasing the distance from veins to stomata, thereby improving photosynthetic efficiency by increasing leaf-level water use efficiency while limiting water loss (Brodribb *et al*., 2007; Bertolino *et al*., 2019). This allows for more carbon to be assimilated per unit of water used by the plant. Because smaller stomata can typically close more quickly due to faster ion fluxes, stomatal conductance can likewise change quickly (Bertolino *et al*., 2019; Drake *et al*., 2013). This behavior allows plants to reduce transpiration in response to drying while still allowing them to rapidly “gear up” when water is available (Sack and Scoffoni, 2013).

Studies of leaf trait plasticity in response to drought have also found a general tendency toward increasingly large trait changes across increasing stress severity. More specifically, decreases in stomatal size and increases in stomatal density and VLA are generally more pronounced as drought severity increases (Lei *et al*., 2018; Wang *et al*., 2018; Stojnić *et al*., 2015; Bresta *et al*., 2018). For instance, there was an increase in stomatal and vein density with increased drought severity in cotton (Lei *et al*., 2018). Similarly, more severe drought resulted in increased stomatal and vein density along with decreased stomatal size in barley (Bresta *et al*., 2018). These observations suggest a potential role for leaf trait plasticity in mitigating the effects of drought, suggesting that the modification of leaf traits might be a reasonable strategy for breeding plants that are more drought resilient. There is, however, a paucity of research that explicitly explores the association of variation in both stomatal and vein traits with performance across multiple levels of drought severity. Here we investigate patterns of leaf trait variation, plasticity, and overall performance in response to varying levels of drought stress in cultivated sunflower (*Helianthus annuus* L.).

Cultivated sunflower is one of the world’s most important oilseed crops (FAO, 2018) and an important source of confectionery seeds and ornamental flowers. Because it is often grown on non-irrigated land, sunflower productivity can be largely dependent on natural patterns of precipitation (Meyer, 1999). While sunflower roots deeply and is able to avoid drought once established (Connor and Sadras, 1992), drought remains a major yield-limiting factor across much of the range of production (Hussain *et al*., 2018). As such, the investigation of drought tolerance and associated traits remains a vital avenue of research. Considering the role that leaf traits play in plant-water relations (Sack and Scoffoni, 2013; Buckley, 2019), they are a potentially important consideration when developing plants that can maintain performance under drought. Previous research in sunflower has shown considerable genetic variability in leaf anatomical traits (Earley *et al*., 2022) as well as substantial phenotypic plasticity in their response to environmental challenges including light- and nutrient-limitation (Wang *et al*., 2020), but little is known about the role of such traits in the response to drought in sunflower. The focus of the present study is on the impact of varying levels of water limitation on leaf anatomical traits and their relationship with overall growth/performance. More specifically, we sought to: (1) evaluate patterns of leaf trait variation and covariation within and across varying levels of water availability; (2) quantify trait plasticity in response to water limitation; and (3) determine the relationship between observed trait variation and plasticity and performance across varying levels of drought stress.

## MATERIALS AND METHODS

### Experimental design

In the summer of 2019, four inbred lines from the sunflower association mapping (SAM) population (Mandel *et al*., 2011) were grown in the Botany Greenhouses at the University of Georgia under a range of watering treatments. These lines correspond to RHA 436 (SAM 33), RHA 364 (SAM 61), INRA line SF 076 (SAM 273), and INRA line SF 075 (SAM 282). Thes lines were chosen to cover a range of stomatal densities based on observations from Earley *et al*. (2022). The experimental design, which included four treatments with four replicates of each genotype in each treatment (N = 4 x 4 x 4 = 64 total individuals), were grown in the greenhouse in a randomized block design. Following germination, all plants were grown for one week in seedling trays to allow for establishment before being transplanted into 7.6 L plastic pots (HPP200; Haviland Plastic Products, Haviland, OH) filled with a 3:1 mixture of sand and a calcined clay mixture (Turface MVP, Turface Athletics). Greenhouse temperature generally ranged from 82-84°F during the day and 72-74°F at night. Individual pots were fertilized with 60 g Osmocote Plus (15-12-9 NPK; Scotts Miracle-Gro, Marysville, OH) and 15 mL each of gypsum (Performance Minerals Corporation, Birmingham, 136 AL) and lime (Austinville Limestone, Austinville, VA) powders for supplemental Ca^2+^. Pots were randomly assigned to one of four treatments: a well-watered control treatment and three drought treatments of varying severity. These treatments were implemented based on the weight of each pot when fully watered. The well-watered control treatment was re-watered daily to 100% of pot capacity while the drought treatments were re-watered to 60%, 40%, and 20% of their capacity based on pot weight (Figure S1). This was done by weighing each pot (with substrate) when fully dry (i.e., substrate before watering and after allowing to sit for three days in the greenhouse) and fully saturated (i.e., pots watered thoroughly and weighed once no longer dripping) to estimate the amount of water required to fully saturate the substrate and determine a target weight for each pot. For context, when converted to gravimetric water content (see below), these treatments correspond to an average of 23.6%, 14.1%, 9.2%, and 4.7% gravimetric water content for the well-watered, mild, moderate, and severe treatments, respectively.

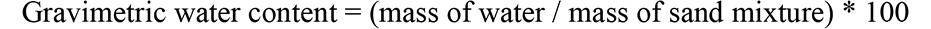

Hereafter, the four treatments will be referred to as the well-watered (100%), mild (60%), moderate (40%), and severe (20%) drought treatments. Following transplantation and initial saturation of the soil, the plants were allowed to acclimate for three days before the treatments commenced. Once the treatments started, pots were re-weighed daily between 9:00-11:00 AM using a postage scale (PS-IN202; Prime Scales, Chino, CA) and allowed to dry to their target levels. Generally, pots for each treatment reached target levels around the same time. Once they dipped below their target weight, pots were individually re-watered to bring them back to their target weight. Note that, while the plants in the severe treatment experienced the most severe drought, most did not reach their target level before the end of the experiment (Figure S1). The drought treatments were maintained for 24 days. Throughout the experiment, plant weight was considered negligible relative to pot/soil weight and was thus not considered when weighing. All individuals were maintained in this way until they reached five weeks of age, at which point they were harvested.

At harvest, traits were measured as described in Earley *et al*. (2022). Briefly, on harvest day, two of the most recent fully expanded leaves (MRFELs) were collected for analysis: one for estimation of LMA and one for anatomical characterization. Prior to removal, chlorophyll content index was measured (Apogee MC-100) on the latter (“anatomy”) leaf for each plant. Following harvest, the LMA leaf was scanned on a flatbed scanner to determine leaf area, dried and weighed to determine the dry biomass of the lamina and midrib (excluding the petiole), and LMA was calculated as: dry mass/unit area (g/m^2^). All remaining biomass was likewise collected, dried, and weighed to determine root, stem, bud, and leaf biomass, which could be summed into total and aboveground biomass. These values were then used to calculate stem, leaf, bud, and root mass fractions by dividing each by the total biomass. Note that bud biomass was not used in further analyses because only a subset of plants had produced buds at the time of sampling.

The anatomy leaf was cut in half lengthwise along the midrib and one half was dried and stored. This half was rehydrated overnight in water at a later date and the adaxial and abaxial (hereafter top and bottom) surfaces were pressed into dental putty (President Dental Putty; Coltène/Whaledent Inc., Cuyahoga Falls, OH) to produce an impression of the epidermis that could be used to visualize and analyze stomatal traits following the general methods of Weyers and Johansen (1985). Nail polish was applied to each impression and lifted off with tape and imaged; this was done separately for the top and bottom surfaces of each leaf (Hilu and Randall, 1984; Weyers and Johansen, 1985). Traits of interest included: stomatal density sum (top + bottom), stomatal length, stomatal pore length, and guard cell width (Earley *et al*. 2022). The total number of stomata was also estimated by multiplying stomatal density sum by leaf area. Imaging for stomatal density and size involved taking images of both the top and bottom impressions for stomatal density and size measurements at 5x and 100x, respectively. Maximum stomatal conductance (*g_smax_*), the theoretical maximum rate of gas exchange if all stomata were fully open, was calculated based on stomatal density and size measurements. This was used instead of directly measuring *g_s_* since it was not feasible to measure directly for all plants. Maximum stomatal conductance was calculated using the approach of (Dow *et al*., 2014).

The second half of the anatomy leaf was stored in Formalin-Acetic Acid-Alcohol (FAA) fixative for imaging and analysis of vein traits. Each sample was cleared, stained, and imaged as described in Earley *et al*. (2022). This included both scanning on a flatbed scanner and imaging four different fields of view under a microscope at 5x. Second order vein length was measured from the scanned images by manually tracing veins that branched off of the midrib (i.e., the primary vein). Major vein length was then determined by adding the length of the midrib to the sum of the second order vein lengths. Minor vein length was measured from the microscope images using a deep neural network as described in Earley *et al*. (2022). Finally, a composite trait of stomata number per vein length was calculated as SV = average [top + bottom] stomatal density/VLA (Zhao *et al*., 2017).

### Data analysis

All data analyses were conducted using R v3.4.3 (R Core Team, 2013) in R Studio v1.3.1093 (RStudio Team, 2015). A two-way ANOVA with genotype and block as the main effects was used to test for variation among genotypes and treatments and to calculate estimated marginal trait means (emmeans; Lenth, 2020) for each genotype after removing block effects. These values were then used to calculate the relative distance plasticity index (RDPI) of each genotype for each trait and treatment. RDPI, which was calculated as described by Valladares *et al*. (2006), but modified to range from -1 to -1 to show the direction of change, allows for cross-treatment comparisons and is strongly correlated with the majority of other measures of phenotypic plasticity. It was calculated for each trait and all treatments versus the well-watered control, as follows:

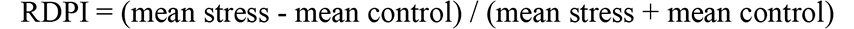

RDPI was also calculated at the pot (i.e., individual plant) level using the same formula except it was calculated for each genotype/treatment combination separately for each block. Therefore, the same control value for each genotype within each block was used in each RDPI calculation. This resulted in 48 individual RDPI measurements.

Pot-level data were used to estimate bivariate trait correlations and to create correlation matrices per treatment using the R package *corrplot* (Wei and Simko 2017). Mantel tests were performed using correlation matrix data and the R package *vegan* to compare correlation matrices and test for differences across treatments (Oksanen *et al*., 2013). A principal component analysis (PCA) of traits by treatment was conducted using the function prcomp() and the package *ggfortify* (Tang *et al*., 2016; Horikoshi and Tang 2018) to visualize the trait-trait correlations. Differences between treatments were investigated using Hotellings *t^2^* test on the first two principal components. All graphs were made using the R package *ggplot2* (Wickham, 2016). Correlation matrices and PC analyses were also conducted using RDPI values for traits within each level of water limitation compared to control to view how plasticity varied across treatments.

A linear mixed effects regression model using the package *brms* (Bürkner, 2017; Bürkner, 2018; Bürkner, 2021) was used to evaluate the relationship between trait variation and variation in performance within each treatment. Given the highly correlated nature of many of the traits in our dataset, we selected four leaf traits that reflect important axes of leaf trait variation in cultivated sunflower (see Earley *et al*., 2022). Traits of interest included: stomatal length and density (important structural traits that influence gas exchange; Lawson and Blatt, 2014), VLA (a venation trait related to leaf water transport; Sack and Scoffoni, 2013), and LMA (which reflects the investment in biomass and leaf construction; Wright *et al*., 2004). We centered and scaled all traits to have a mean of zero and standard deviation of one. Due to a few extreme LMA values, we log-transformed LMA values before centering and scaling to improve model fit. The model included the four leaf traits and the interaction of each with stress level as main effects. Block and genotype ID were included as random intercepts in the model. Default priors were used (Bürkner, 2017; Bürkner, 2018; Bürkner, 2021) with a step size of 0.99, four chains with 2000 iterations (with 1000 iterations for a warm-up), and no thinning for a total of 4000 samples from the posterior. The model was evaluated for efficient mixing. No Rhat values greater than 1.00 were reported and there were no divergent transitions. The treatment level-specific coefficients relating each leaf trait to total biomass are reported below, as are the Bayesian marginal *R*^2^ estimates (Gelman *et al*., 2019) which estimates the variance explained by the fixed effects in the model.

A similar model was fit to evaluate the association between the plasticity of a trait (i.e., the RDPI value, compared to control) and the RDPI in biomass to investigate the association between shifts in trait values and changes in performance across treatments. The only difference in this model is that we did not include the interaction with treatment level, and instead included treatment as a random intercept to account for multiple comparisons (i.e., each trait RDPI in each of the three stress treatments was compared with the same control). We used the same model parameters and report no Rhat values greater than 1.00 and no divergent transitions.

## RESULTS

### Leaf trait variation and covariation within and across treatments

The median, minimum, and maximum values for all traits, along with a summary of the ANOVA results for each, are listed in Table 1. See also Figure 1 for a visual presentation of data for four representative traits. Tukey’s HSD test results for all traits across levels of treatment severity are presented in Table S1. In general terms, biomass decreased as severity increased while stomatal and vein density increased with increasing severity. In contrast, *g_smax_* remained largely constant across all but the most severe treatment. Significant genotypic effects were detected for a subset of leaf anatomical as well as higher-level (i.e., whole plant) traits. This included traits related to guard cell width, several traits related to venation, leaf size, total and aboveground biomass, and multiple mass fractions. Significant treatment effects were found for nearly all traits with the exception of guard cell width, SV, chlorophyll, and bud mass fraction. There was, however, minimal evidence of genotype-by-treatment interactions; exceptions to this included significant interactions for leaf area, midrib density, total biomass, and total stomata.

**Figure 1:**
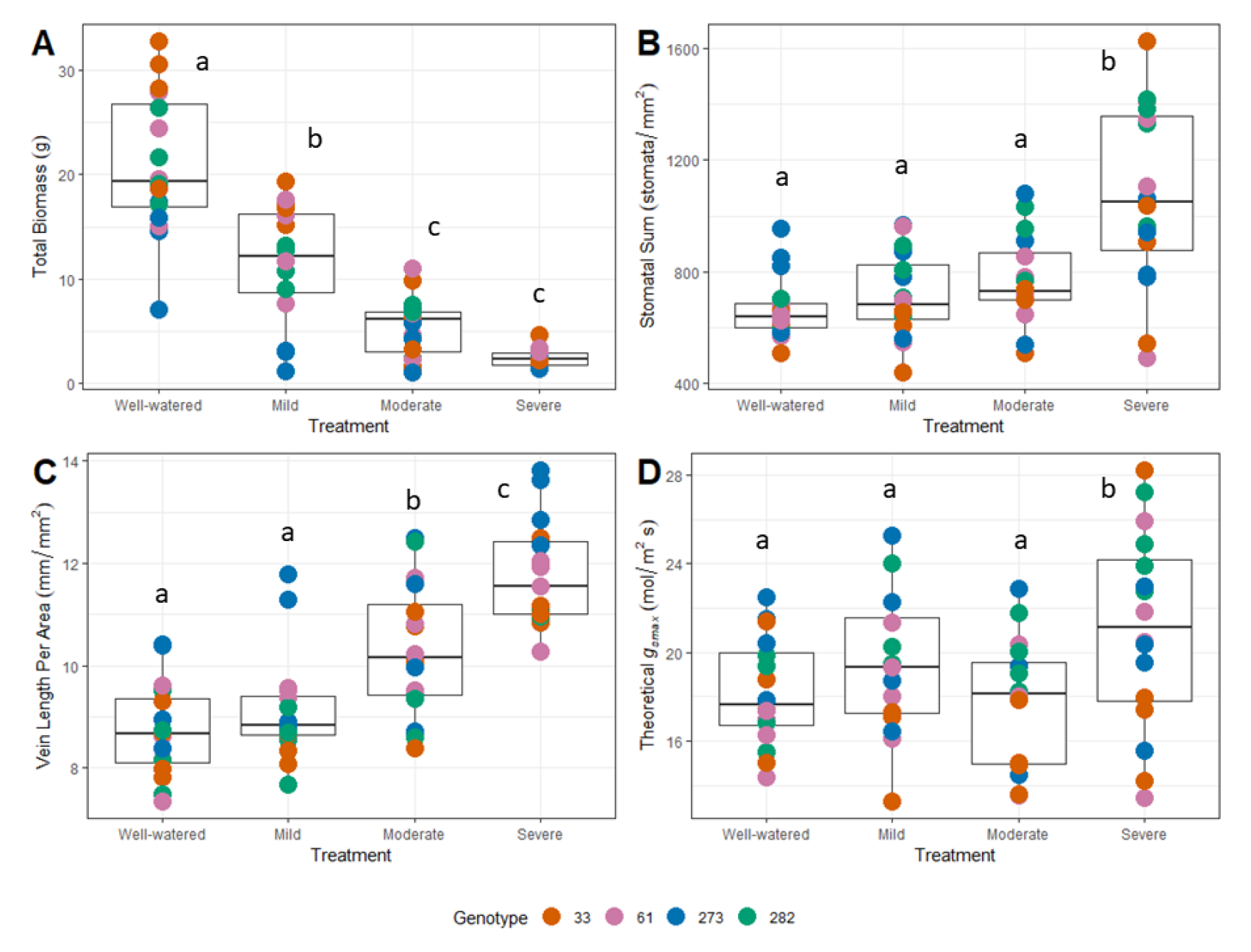
Boxplots of variation in representative traits across treatments. Dots are colored based on genotype, which are listed by line number in the key at the bottom. Traits of interest include: (A) total biomass; (B) stomata density sum; (C) vein length per area; and (D) theoretical *g_smax_*. Letters indicate significant differences across treatments based on Tukey’s HSD test.

**Table 1:**
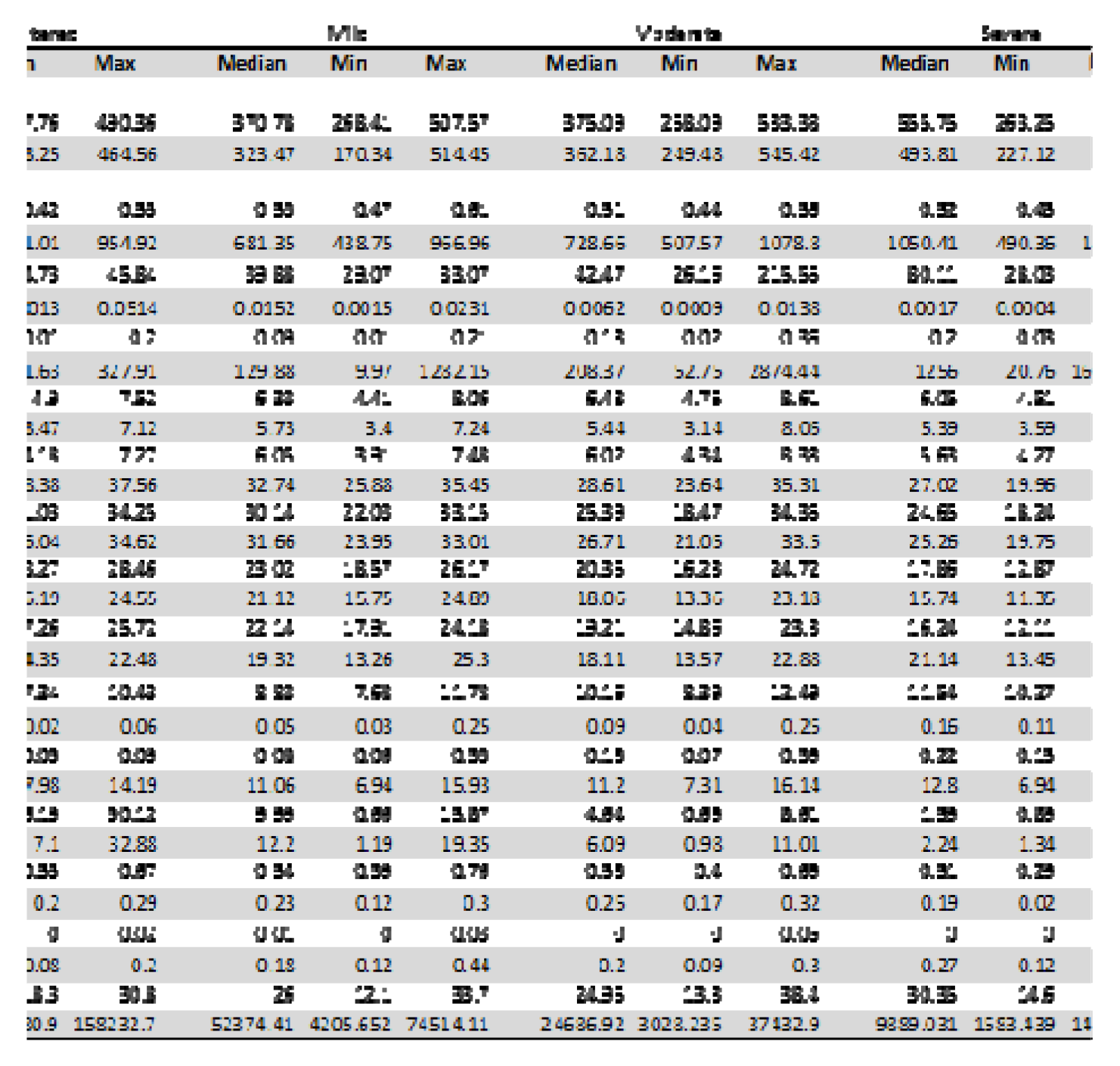
List of all traits measured along with the median and range of trait values. Significance for genotypic effects (***P ≤ 0.001, **P ≤ 0.01, * P ≤ 0.05, **#** P ≤ 0.1, ns = not significant) and adjusted R2 from the model are also presented. SD = stomatal density; LMA = leaf mass per area; MF = mass fraction; GCW = guard cell width; SL = stomatal length; PL = stomatal pore length; g_smax_ = theoretical maximum stomatal conductance; VLA = vein length per area; SV = stomata per vein length; AG = aboveground; Bio = biomass.

All bivariate trait correlations are presented in correlations matrices for each treatment (Figure S2). Mantel tests revealed that none of the matrices were significantly different from one another (i.e., the null hypothesis of a lack of correlation between matrices was rejected [*P < 0.001*] for all comparisons) suggesting that the overall correlation structure does not change appreciably across treatments. Multivariate trait relationships were analyzed via PCA to determine the major axes of variation (Figure 2). In the case of traits with values for the top and bottom of leaves, which are strongly correlated, averages were used to simplify the analysis and interpretation. Because overall size differences across treatments would largely obscure other trait relationships, this analysis was based on putatively size-independent traits (i.e., traits that lacked a significant correlation with size-related measures), with leaf area and both aboveground and total biomass being excluded. Second VLA was also removed due to its redundancy with major VLA. For a full PCA including all measured traits see Figure S3. For the analysis of size-independent traits, the top three principal components accounted for 67.5% of the observed trait variation (Figure 2; Figure S4) with PC1 explaining 35.2%, PC2 explaining 16.6%, and PC3 explaining 15.7%. The top three traits contributing to each major axes were: PC1 - stomatal density sum, stomata length (SL_Avg), and stomatal pore length average (PL_Avg); PC2 - chlorophyll content, stomata per vein length (SV), and theoretical *g_smax_*; and PC3 - leaf mass fraction, root mass fraction, and LMA (Table 2). Consistent with the observations of Earley *et al*. (2022), the first three PCs seem to be heavily influenced by traits relating gas exchange, hydraulics, and leaf construction.

**Figure 2:**
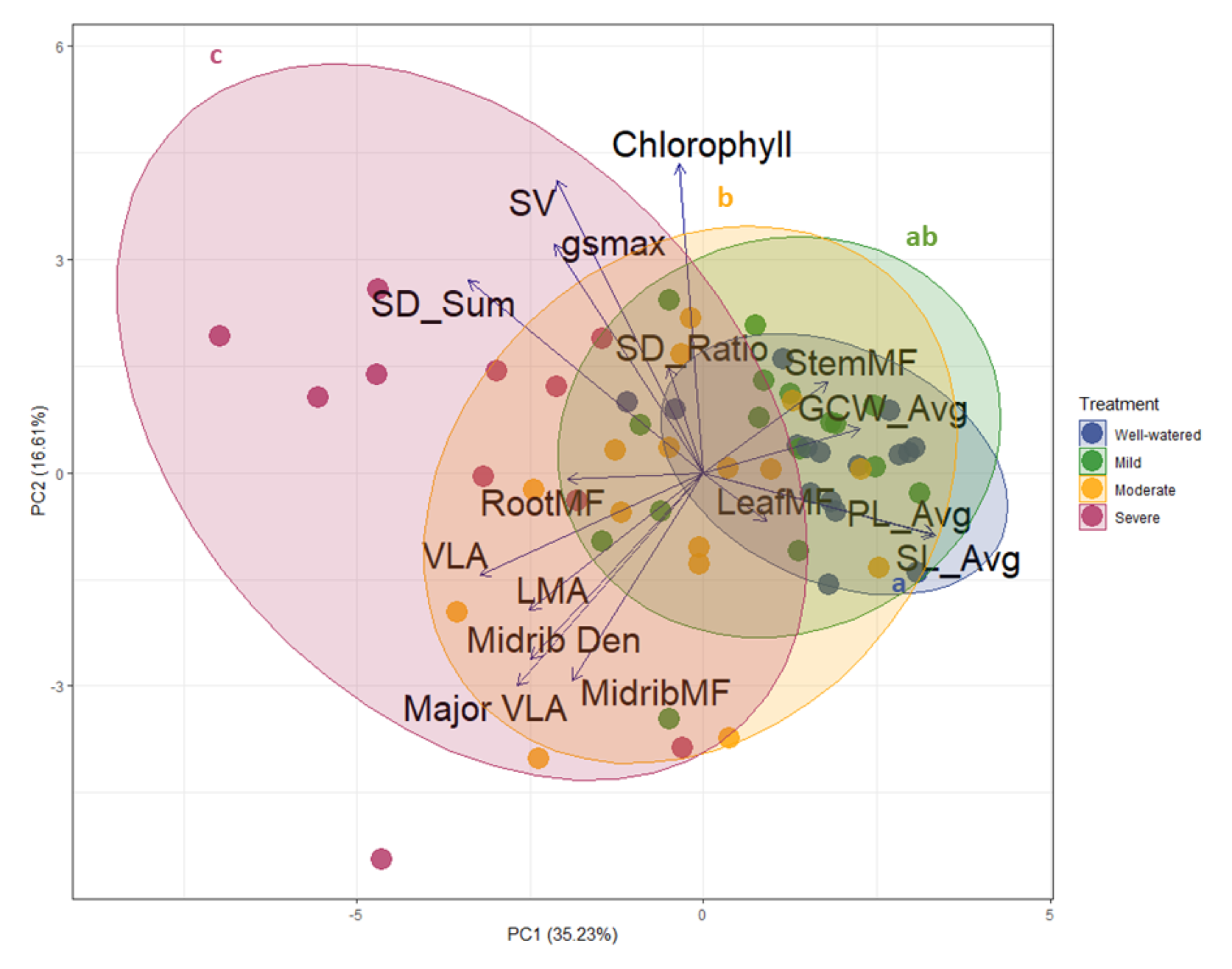
Principal component analysis (PCA) of putatively size-independent traits. Traits names that include _Avg (stomatal length [SL], pore length [PL], guard cell width [GCW]) are averages of the values from the top and bottom of the leaf. SD_Sum is the sum of stomatal density from the top and bottom of the leaf. Colors indicate treatment and trait abbreviations are as defined in Table 1. Lower-case letters associated with each of the colored ellipses indicate the results of Hotelling’s t^2^ tests for differences between treatments. P-values were adjusted for multiple comparisons using a Bonferroni correction.

**Table 2:**
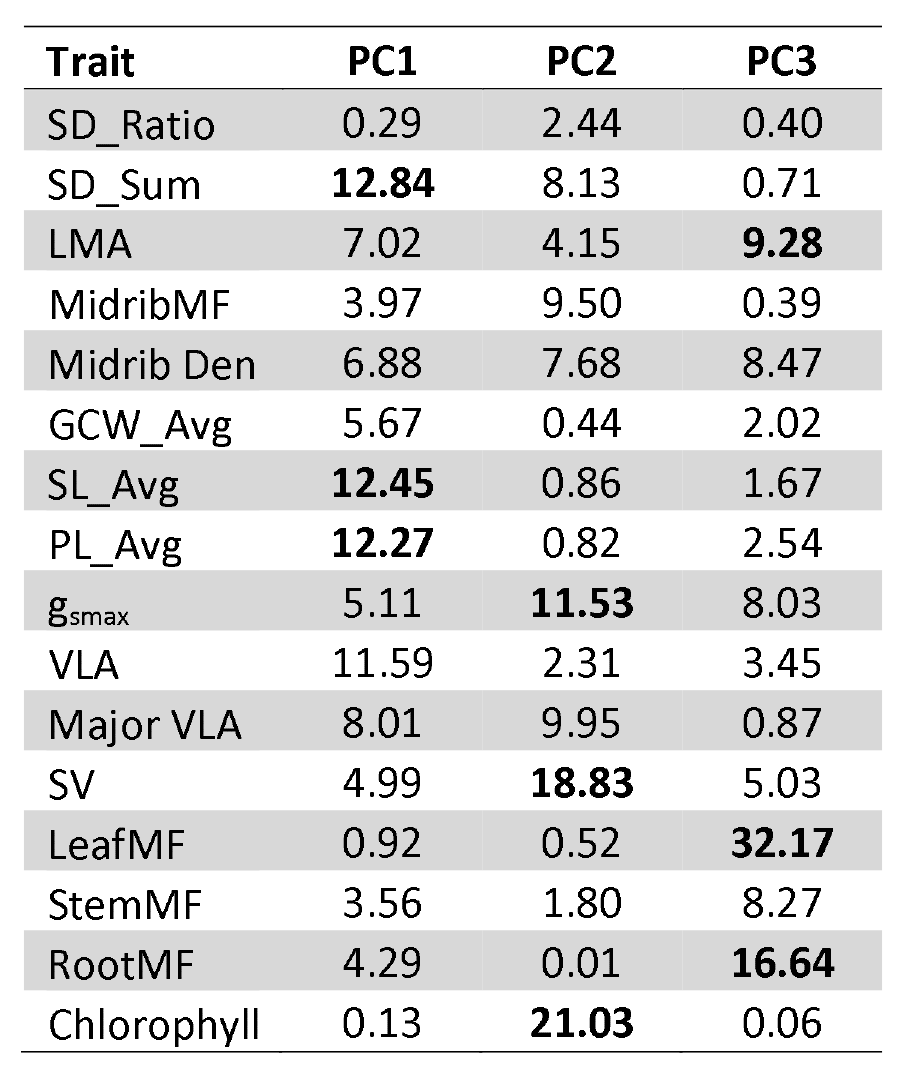
Trait loadings (percentage of trait variation explained by each trait in the associated principal component [PC]) for each of the first three PCs. The top three traits per PC are highlighted in bold. Trait abbreviations follow Table 1.

| Trait | PC1 | PC2 | PC3 |
| --- | --- | --- | --- |
| SD_Ratio | 0.29 | 2.44 | 0.40 |
| SD_Sum | **12.84** | 8.13 | 0.71 |
| LMA | 7.02 | 4.15 | **9.28** |
| MidribMF | 3.97 | 9.50 | 0.39 |
| Midrib Den | 6.88 | 7.68 | 8.47 |
| GCW_Avg | 5.67 | 0.44 | 2.02 |
| SL_Avg | **12.45** | 0.86 | 1.67 |
| PL_Avg | **12.27** | 0.82 | 2.54 |
| $g_{smax}$ | 5.11 | **11.53** | 8.03 |
| VLA | 11.59 | 2.31 | 3.45 |
| Major VLA | 8.01 | 9.95 | 0.87 |
| SV | 4.99 | **18.83** | 5.03 |
| LeafMF | 0.92 | 0.52 | **32.17** |
| StemMF | 3.56 | 1.80 | 8.27 |
| RootMF | 4.29 | 0.01 | **16.64** |
| Chlorophyll | 0.13 | **21.03** | 0.06 |

### Trait plasticity in response to water limitation

When calculated using a relative distance plasticity index (RDPI; Valladares *et al*., 2006), estimates of trait plasticity can range from -1 to 1 with positive or negative values reflecting increases or decreases in trait values in response to the treatment, and 0 representing no plasticity. Overall, trait plasticity (as compared to control) increased with increasing drought severity for nearly all traits and was consistent across all genotypes (Figure 3). In general, smaller-scale leaf anatomical traits such as stomatal size and density and VLA exhibited smaller plasticity estimates than larger-scale traits related to construction and growth, such as second/major VLA, leaf area, and biomass.

**Figure 3:**
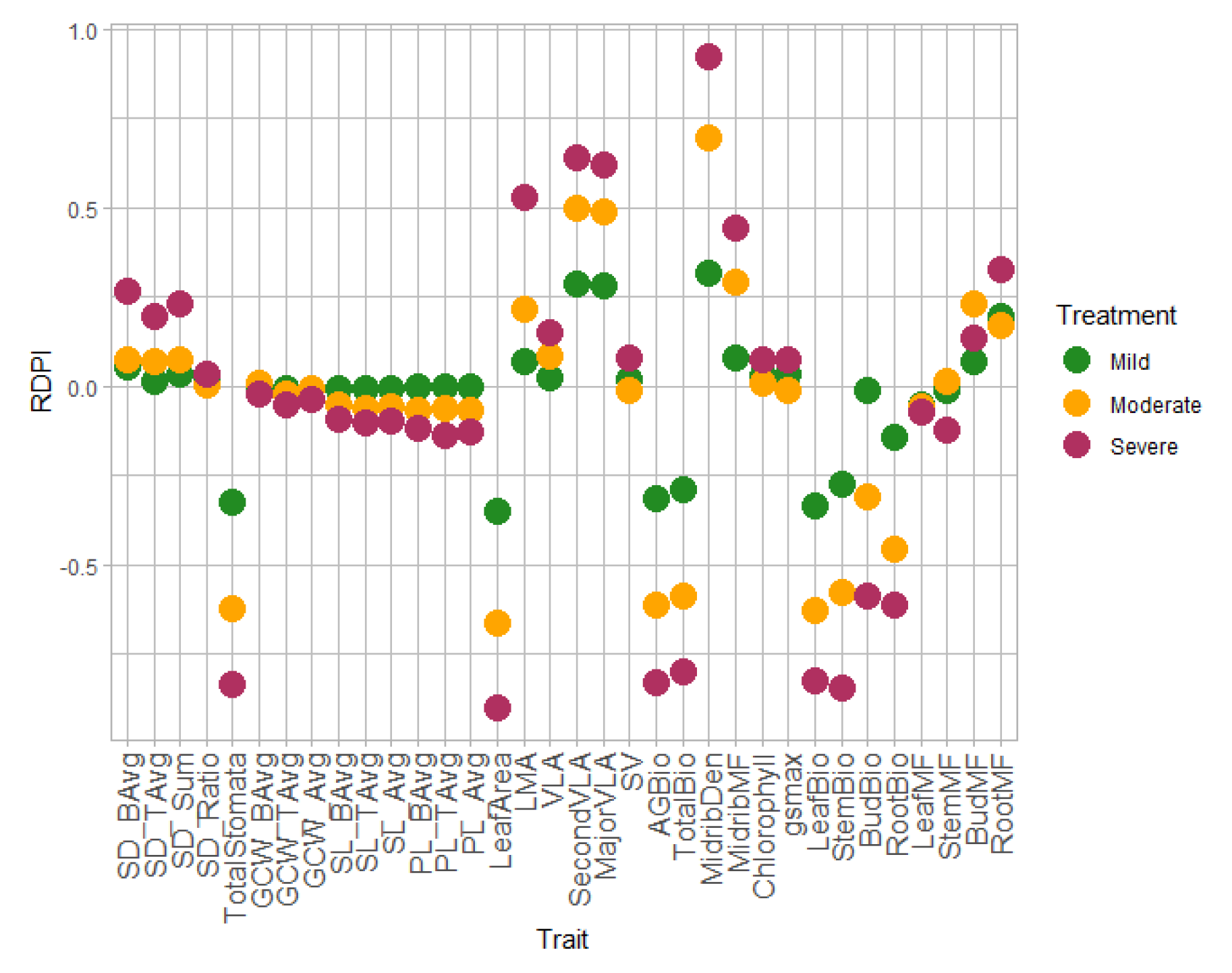
Relative distance plasticity index (RDPI) plot for each trait and treatment. Points indicate the average estimated marginal mean for each trait for each treatment. RDPI was calculated as each treatment versus the well-watered control. Trait abbreviations follow Table 1.

Analysis of multivariate relationships for the RDPI values of putatively size-independent traits were analyzed via PCA. The top three PCs account for 68.2% of the observed variation in plasticity values with PC1 explaining 29.0%, PC2 explaining 22.2%, and PC3 explaining 17.0% (Figure 4). The top three plasticity values contributing to each major axes corresponded to the following traits: PC1 - stomatal density sum, stomata length (SL_Avg), and stomatal pore length average (PL_Avg); PC2 - chlorophyll content, stomata per vein length (SV), and midrib mass fraction; PC3 - theoretical *g_smax_,* leaf mass fraction, and root mass fraction (Table 3). As seen for the trait PCA, the first three PCs are heavily influenced by plasticity in traits relating to hydraulics, gas exchange, and construction (Table 3). For a PCA of the RDPI for all traits see Figure S5.

**Figure 4:**
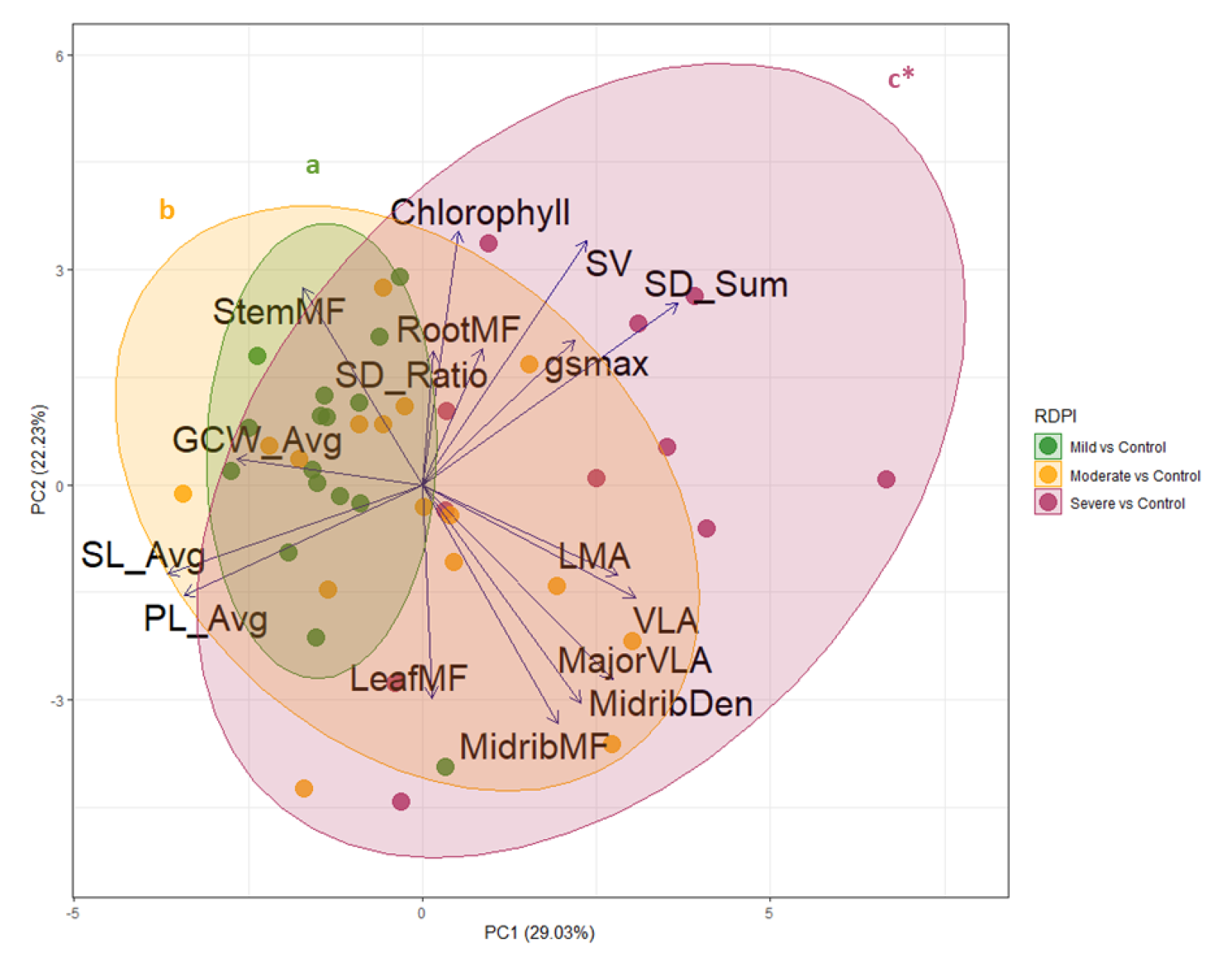
Principal component analysis (PCA) of trait plasticity values (estimated as RDPI) for each treatment compared to control. Trait abbreviations follow Table 1. Lower-case letters associated with each of the colored ellipses indicate the results of Hotelling’s t^2^ tests for differences between treatments. P-values were adjusted for multiple comparisons using a Bonferroni correction. *Severe vs. control is marginally significant following correction for multiple comparisons.

**Table 3:** Trait loadings (percentage of trait variation explained by each trait in the associated principal component [PC]) for each of the first three PCs of plasticity (RDPI) values. The top three traits per PC are highlighted in bold. Trait abbreviations follow Table 1.

| Trait | PC1 | PC2 | PC3 |
| --- | --- | --- | --- |
| SD_Ratio | 0.03 | 3.68 | 3.92 |
| SD_Sum | **14.37** | 6.79 | 1.58 |
| LMA | 8.41 | 1.69 | 6.17 |
| MidribMF | 4.05 | **11.84** | 2.27 |
| MidribDen | 5.55 | 9.86 | 6.89 |
| GCW_Avg | 7.60 | 0.14 | 7.68 |
| SL_Avg | **14.24** | 1.66 | 5.25 |
| PL_Avg | **12.51** | 2.54 | 5.28 |
| $g_{smax}$ | 5.12 | 4.32 | **14.94** |
| VLA | 9.98 | 2.66 | 7.41 |
| MajorVLA | 7.96 | 7.88 | 2.96 |
| SV | 5.95 | **12.34** | 7.00 |
| LeafMF | 0.02 | 9.44 | **13.25** |
| StemMF | 3.13 | 8.03 | 1.24 |
| RootMF | 0.78 | 3.83 | **8.57** |
| Chlorophyll | 0.28 | **13.31** | 5.60 |

### Relationship between trait variation and/or trait plasticity and performance

Our model using four key traits selected to represent variation in major axes of leaf trait variation (i.e., vein length per area, stomatal length, stomatal density [sum], and LMA) to predict plant performance across environments (measured as total biomass) exhibited strong predictive power with an overall model marginal *R*^2^ = 0.74 (Figure 5A). The treatment-specific estimated associations between each trait and biomass are presented in Figure 5B and the effect of stress on productivity (biomass) is illustrated in Figure 5C. For coefficients and credible intervals, see Table S2. Uncertainty in model estimates was evaluated using posterior credible intervals. Estimates with 95% CIs not overlapping zero were interpreted as having strong support of the association, and those with 80% CIs with moderate support. VLA was negatively associated with biomass under control and mild stress conditions, but positively associated under severe stress (the effect estimated for moderate stress was positive, but the 80% CI overlapped with zero). Stomatal length had no clear association with biomass under control conditions, mild, or severe stress, and a positive association under moderate stress. Stomatal sum had a positive association with biomass under control and moderate stress conditions, but no association under mild and severe stress. Finally, log-LMA had a positive association with biomass under control conditions but no clear association under any of the stress conditions (Figure 5B).

**Figure 5:**
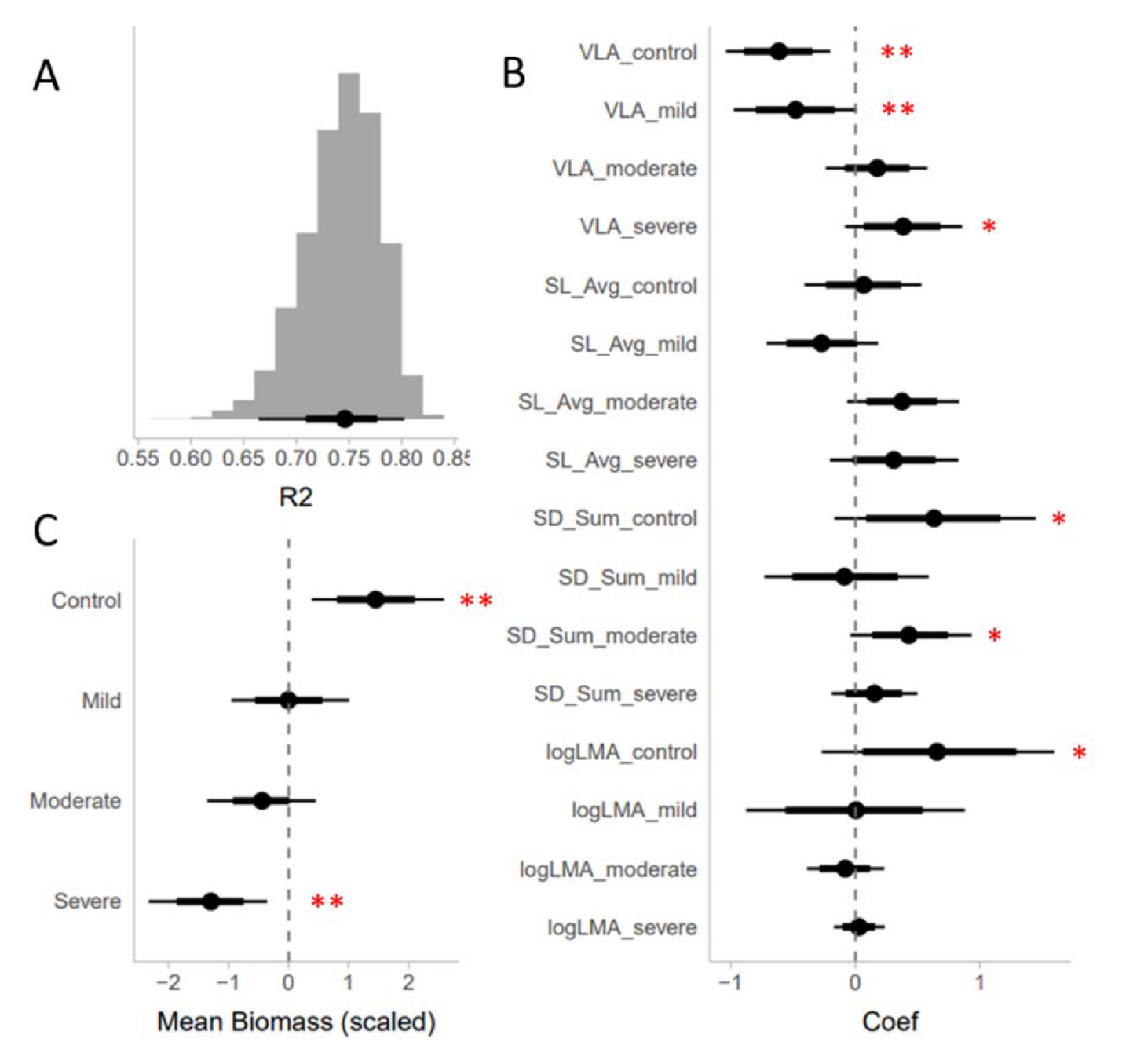
Results of our analysis of performance as a function of variation in VLA, average stomatal length (SL_Avg), stomatal density (SD_sum), and LMA across environments using a linear mixed effects regression model. All output reflects posterior summaries from the model. (A) Estimated Bayesian R2, for fixed effects only (full posterior distribution of the model). (B) Coefficient estimates for each trait/environment combination. LMA was log-transformed to improve model fit and all traits were scaled to a mean of zero and standard deviation of one. (C) Mean biomass for each treatment scaled to a mean of zero and a standard deviation of one. Black dot indicates the estimated mean value. The heavier bars indicate the 80% credible interval for each estimate, and the lighter lines indicate the 95% credible interval. Estimates with 95% CIs not overlapping zero were interpreted as having strong support (**), and those with 80% Cis with moderate support (*).Plots of (C) VLA vs total biomass and (D) stomatal density sum vs total biomass. Dots represent individuals and lines are the line of best fit per treatment.

The results of the model using plasticity estimates (i.e., RDPI values) for the same traits as above (i.e., VLA, stomatal length, stomatal density, and LMA) revealed the ability to predict plasticity in plant performance with a marginal model *R*^2^ = 0.25 (Figure 6A). Individual coefficient estimates are presented in Figure 6B. For coefficients and credible intervals see Table S3. The RDPI of both VLA and stomatal density were associated with changes in biomass, but in different directions (negatively and positively, respectively). The resulting pattern is one in which higher VLA RDPI is associated with more extreme decreases in biomass (and vice versa), while higher stomatal density RDPI is associated with less extreme decreases in biomass (Figures 6C and 6D). In contrast, the RDPI of stomatal length and LMA had no association with the change in biomass.

**Figure 6:**
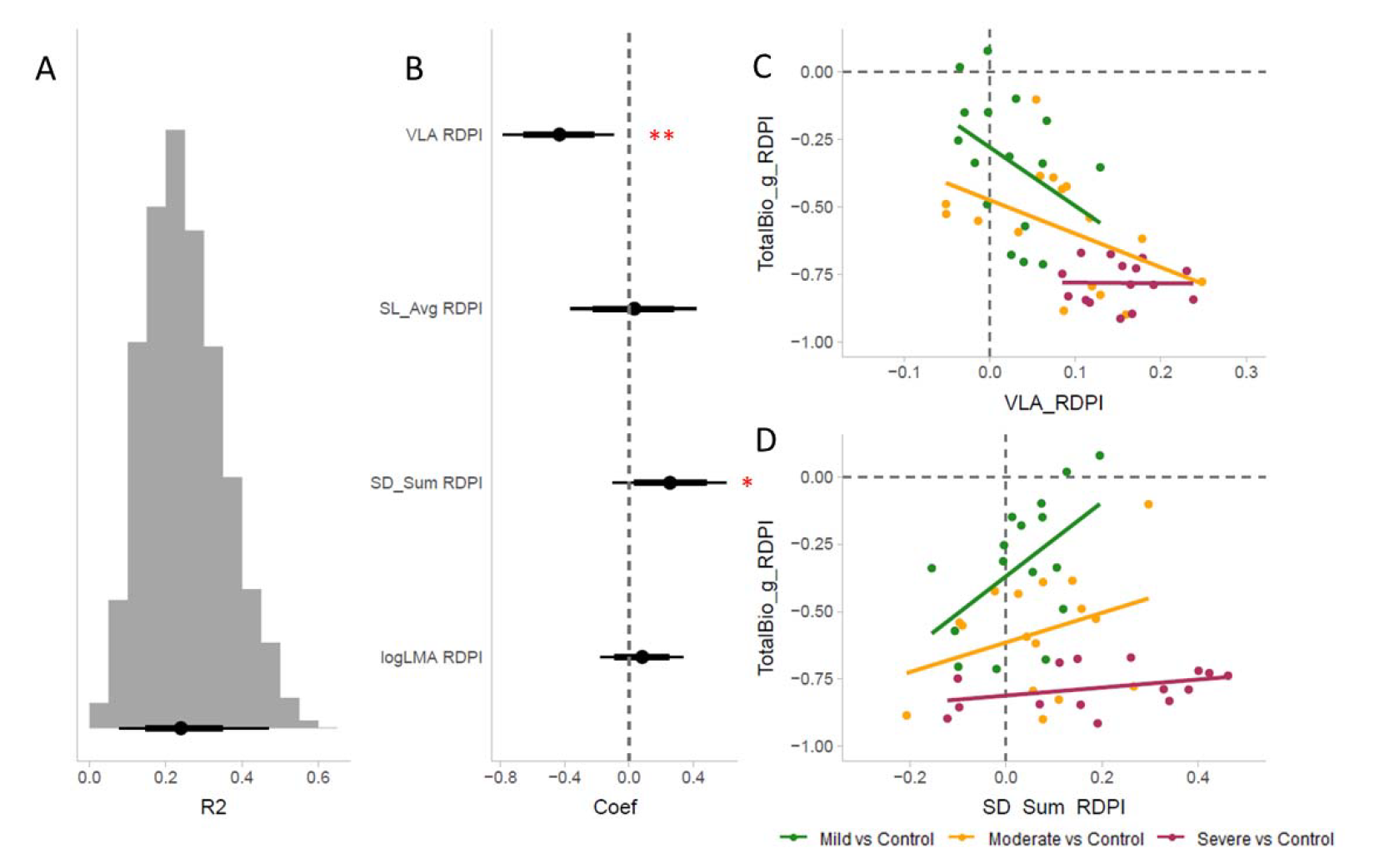
Results of our analysis of plasticity in performance as a function of variation in plasticity (RDPI) of VLA, average stomatal length (SL_Avg), stomatal density sum (SD_sum), and LMA using a linear mixed effects regression model. All output reflects posterior summaries from the model. (A) Estimated Bayesian R2, for fixed effects only (full posterior distribution of the model). (B) Coefficient estimates for each trait. LMA was log-transformed, and all traits were scaled to a mean of zero and standard deviation of one. Black dot indicates the estimated mean value. The heavier bars indicate the 80% credible interval for each estimate, and the lighter lines indicate the 95% credible interval. Estimates with 95% CIs not overlapping zero were interpreted as having strong support (**), and those with 80% CIs with moderate support (*).

## DISCUSSION

Drought is a major challenge that reduces plant growth and limits crop productivity worldwide. Stomatal and vein traits play an important role in plant-water relations and are thus of interest in the context of drought tolerance. Here, we investigated patterns of leaf trait covariation within and across treatments in cultivated sunflower. We also sought to quantify leaf trait plasticity in response to water limitation and determine the extent to which trait variation and/or plasticity predicts performance across treatments.

### Leaf trait variation and covariation within and across treatments

The existence of significant genotypic effects for traits related to leaf area, stomatal morphology, venation, and growth/allocation suggests the occurrence of genetic differentiation for a wide range of traits across cultivated sunflower lines. Within treatments, traits exhibited many of the predicted relationships based on the literature and their known roles in plant-water relations (Sack and Scoffoni, 2013; Doheny-Adams *et al*., 2012; Shahinnia *et al*., 2016; Figure S2). For example, stomatal density and size were negatively correlated while stomatal density and VLA were positively correlated (Figure S2). The former relationship indicates that the total area allocated to stomata affects total stomatal conductance and thus total photosynthesis (Harrison *et*□*al*.□2019; Shahinnia *et*□*al*. 2016). The latter relationship likely represents a balance between stomata and veins such that water use and carbon acquisition are optimized (Carins Murphy *et*□*al*.L2014; Brodribb *et*□*al*.□2007; Sack and Scoffoni□2013).

Notably, there was minimal evidence of trait variation scaling with size within treatments. The primary exception was that second and major VLA are negatively correlated with various aspects of plant size, whereas leaf size is negatively correlated with second and major VLA. This makes sense, as larger plants typically had larger leaves, and major veins are formed early in leaf development before being pushed apart as leaf expansion accelerates (Sack and Scoffoni,L2013). Minor veins are expected to show no such relationship (as seen here) because they can be initiated throughout leaf development. While total stomatal count was positively correlated with leaf size, no other stomatal or vein traits were associated with leaf size, indicating that correlations between combinations of those traits exist independent of leaf size as opposed to being a byproduct of allometric scaling.

Interestingly, the structures of the correlation matrices were largely conserved across treatments, suggesting that the observed trait relationships are quite robust to environmental perturbations. However, when viewing major axes of trait variation using PCA, differences begin to emerge (Figure 2). For example, the trait space occupied by individuals in the severe stress treatment was significantly different from all other treatments. Similarly, the moderate and well-watered treatments were significantly different from each other, though mild was not detectably different from either of them. Across treatments, individuals separated along major axes of variation corresponding to traits involved in gas exchange, hydraulics, and construction. Interestingly, similar axes of variation were observed in a prior study of a large cultivated sunflower diversity panel (Earley *et al*., 2022) suggesting the existence of a persistent functional relationship amongst these traits that could mediate interactions with the environment.

### Trait plasticity in response to water limitation

As expected, drought stress resulted in decreased biomass production, with the magnitude of the effect increasing with increased stress severity (Figure 1A). This decrease in biomass was presumably due to water limitation reducing the capacity for photosynthesis, thereby slowing or stopping growth (Munns, 2011; Kruger *et al*. 2006). In addition to this overall effect on growth, allocation patterns changed with root mass fraction increasing and stem and leaf mass fractions slightly decreasing (Table 1). For the most part, these allocation changes reflect common responses to drought stress (Eziz *et al*., 2017). By shifting resources into roots, plants are effectively maximizing their ability (given their decreased overall size) to find and acquire water during drought. Given that leaves are a source of water loss, decreases in leaf biomass might also help limit loss during times of limited water availability.

In general terms, we observed an increase in stomatal density and VLA along with a decrease in stomatal size in response to drought (Table 1; Figure 1). Similar phenotypic responses have been documented in response to water limitation in other plant species (e.g., Xu and Zhou 2008; Sun *et al*., 2014) and may play a role in increasing water use efficiency under stress (Xu and Zhou, 2008; Lei *et al*., 2018). These trait shifts also reflect observed patterns of genetic differentiation both within (Xie *et al*., 2022; Dunlap and Stettler, 2001) and among species (de Boer *et al*., 2016; Strobel and Sundberg, 1983), wherein individuals from increasingly dry habitats tend to exhibit smaller, denser stomata with an associated increase in vein density. The plastic responses of leaf anatomical traits thereby appear to parallel putatively adaptive solutions to water limitation that have evolved within and among plant species. Given that smaller stomata are able to close more quickly, this phenotypic response to stress might improve the ability of plants to rapidly adjust stomatal conductance in response to changes in water availability (e.g., Brodribb *et al*., 2007; Bertolino *et al*., 2019; Sack and Scoffoni, 2013) The associated increase in VLA allows for greater hydraulic conductance and the maintenance of hydraulic function is critical for plant survival under drought (Yao *et al*., 2021).

Looking across treatments, we found that the magnitude of plasticity for most traits increased as drought stress intensified (Figure 3). Overall, plasticity (RDPI) estimates were quite large for traits related to biomass production as well as traits related to leaf construction such as second and major VLA, leaf area, and LMA (Figure 3). In contrast, RDPI estimates for leaf anatomical traits such as measures of stomatal size, stomatal density, and VLA were much smaller. Similar trends have been documented in other species when subjected to stress. For example, in a study of plasticity in response to varying light intensities in rice (Chen *et al*., 2021), traits related to leaf morphology and anatomy exhibited less plasticity than growth-related traits. This pattern may result from the coordination of certain trait combinations that need to maintain specific relationships to ensure proper function – e.g., stomatal and vein traits exhibit strong correlations, the maintenance of which helps ensure efficient water use (Brodribb *et al*., 2013). In contrast, growth-related traits are perhaps more free to vary in response to changes in resource availability without impairing function.

When viewed in a multivariate context, the extent of trait plasticity clearly varies across treatments (Figure 4). This is, however, largely due to an increase in the magnitude of response across traits as the severity of the drought treatment increased as opposed to novel trait shifts in response to different stress scenarios. As was the case with the PCA of trait values *per se*, the analysis of trait plasticity (RDPI) values revealed three primary axes of variation. Interestingly, these axes largely matched those observed in the trait-based PCA, with the top contributors of the first three PCs again corresponding to (plastic responses in) traits involved in hydraulics, gas exchange, and construction (Table 3). It thus appears that variation in trait responses across environments is structured in much the same way as trait variation within environments.

### Relationship between trait variation and/or trait plasticity and performance

Our results show that variation in just four size-independent leaf traits is strongly predictive (*R*^2^ = 0.74) of plant performance measured as total biomass production. These four traits (VLA, stomatal length, stomatal density, and LMA) were primarily chosen to reflect the major axes of variation identified in previous work (Earley *et al*., 2022) and documented again herein. This result suggests that these and/or closely related leaf traits are strongly associated with, and potentially impact overall plant performance. This makes logical sense and has been observed in the literature. Given the role that veins, and stomata play in gas exchange and water transport and thus photosynthetic capacity (Sack *et al*., 2013), the observed relationship between these traits and growth performance is perhaps not surprising.

Interestingly, the area of a single leaf was even more strongly predictive of total biomass (*R*^2^ = 0.94). While not a truly size-independent trait, this result points to a single, easy to measure trait that may perform well as a proxy for total biomass in sunflower. As far as we are aware, a similar relationship has not been documented in other plant species. It is important to note that leaf area and VLA are not entirely independent, as the log-transformed values of both traits exhibit a linear relationship (Figure S6). Despite this relationship, we found substantial variation in VLA within each of the four treatments suggesting that this relationship is more complex than just scaling with size. In addition, it has been shown that VLA and stomatal density are coordinated independent of leaf area (Carins Murphy *et al*., 2016).

In terms of trait plasticity, we found that VLA RDPI is negatively associated with biomass RDPI and that stomatal density RDPI is positively associated (Figure 6). In other words, increases in VLA are associated with a more severe drop in biomass under stress. Conversely, increases in stomatal density are associated with a less severe drop in biomass (Figure 6). Given the documented positive correlation between stomatal density and VLA (Earley *et al*. 2022 and herein), the finding that changes in these traits have opposite effects on performance seems counterintuitive. Based on previous work (Earley *et al*. 2022), we would have expected an increase in both to increase performance, but our results here show a positive relationship with stomatal density and a negative relationship with VLA. Ultimately, however, this finding suggests that the covariation of these traits is not strictly constrained and that decoupling them might allow for the exploration of novel phenotypic space that could improve performance under drought.

## ACKNOWLEDGMENTS

We thank Kelly Bettinger, Mike Boyd, Greg Cousins, the rest of the greenhouse staff, and numerous members of the Burke and Donovan labs for help with greenhouse work and harvest. Niki Padgett, Summerlin Courchaine, and Kelly Bettinger provided invaluable assistance with the leaf image processing; Chris Cotter provided the deep learning neural network and Alex Bucksch provided valuable feedback on the automated image analyses; and members of the Burke and Donovan labs provided comments that greatly improved an earlier version of the manuscript. This work was supported by a grant from the NSF Plant Genome Program (IOS- 1444522) to JMB as well as funding from the International Consortium on Sunflower Genomics.

