## Supplemental Figures and Tables for "Leaf traits predict performance under varying levels of drought stress in cultivated sunflower (*Helianthus annuus* L.)"

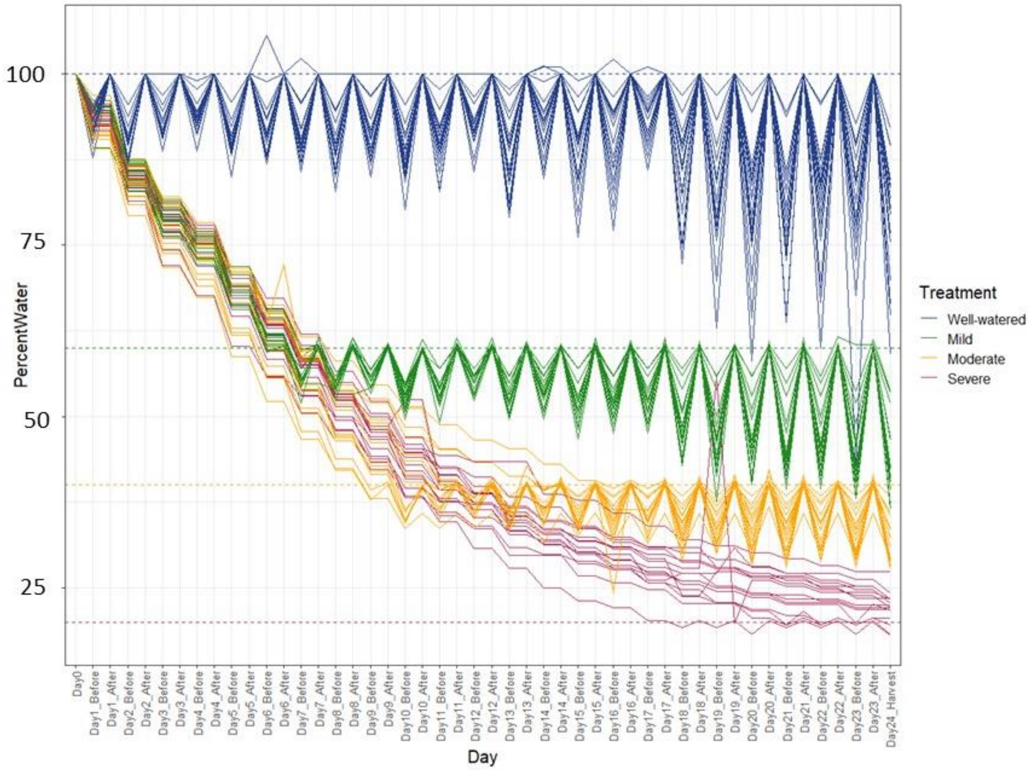


**Figure S1**: Watering data based on percent water compared to field capacity - before and after watering each day. Dotted lines are target levels for each treatment. Well-watered - 100%, mild - 60%, moderate - 40%, severe - 20%.

A

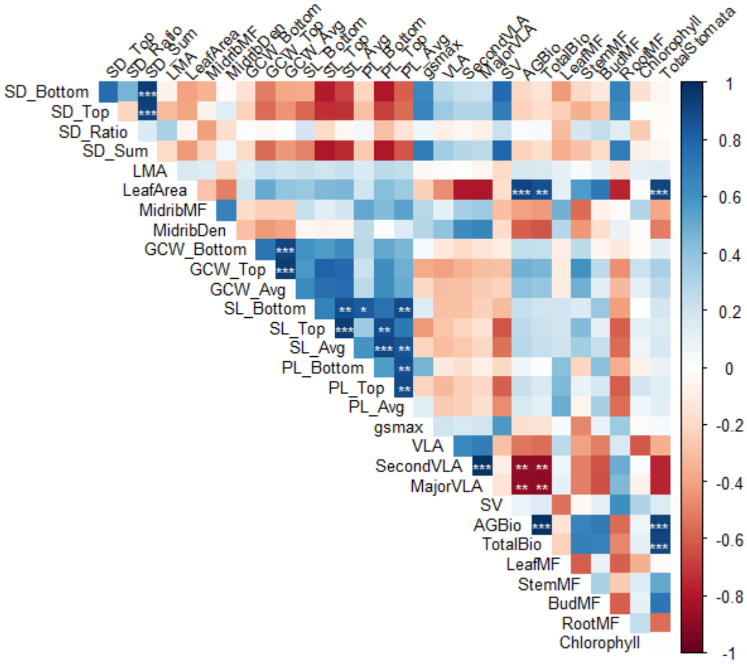
SD_Top
SD_Ratio
SD_Sum
LMA
LeafArea
MidribMF
MidribDen
GCW_Bottom
GCW_Top
GCW_Avg
SL_Bottom
SL_Top
SL_Avg
PL_Bottom
PL_Top
PL_Avg
gsmax
VLA
SecondVLA
MajorVLA
SV
AGBio
TotalBio
LeafMF
StemMF
BudMF
RootMF
Chlorophyll
TotalStomata
SD_Bottom
Chlorophyll
1
0.8
0.6
0.4
0.2
0
-0.2
-0.4
-0.6
-0.8
-1

B

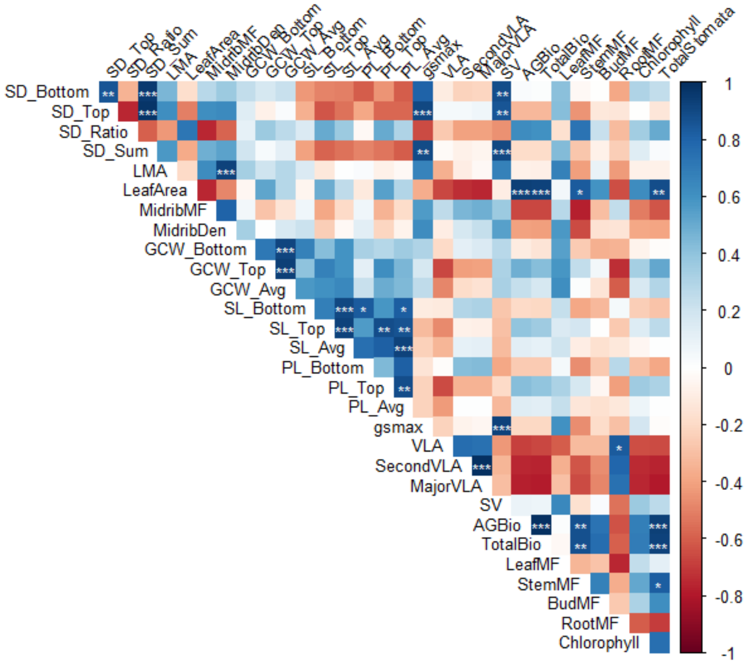
SD_Top
SD_Ratio
SD_Sum
LMA
LeafArea
MidribMF
MidribDen
GCW_Bottom
GCW_Top
GCW_Avg
SL_Bottom
SL_Top
SL_Avg
PL_Bottom
PL_Top
PL_Avg
gsmax
VLA
SecondVLA
MajorVLA
SV
AGBio
TotalBio
LeafMF
StemMF
BudMF
RootMF
Chlorophyll
TotalStomata
SD_Bottom
Chlorophyll
1
0.8
0.6
0.4
0.2
0
-0.2
-0.4
-0.6
-0.8
-1

**C**

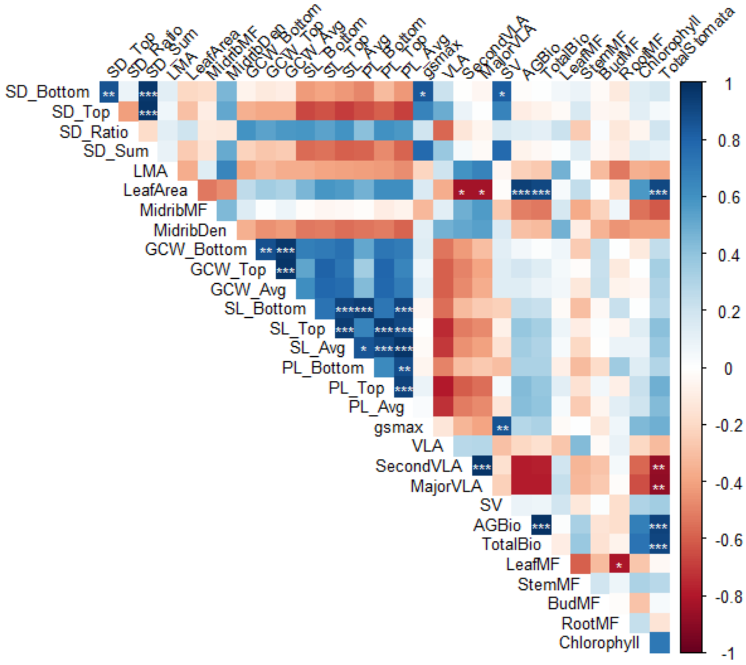

**D**

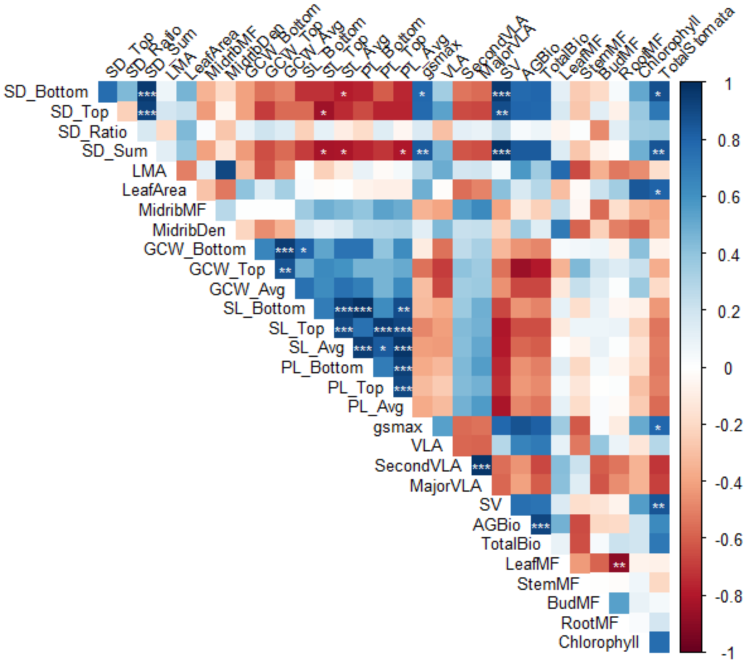

**Figure S2:** Correlation matrices for each treatment (A) well-watered; (B) mild; (C) moderate; (D) severe Values were calculated using pot level data and significance tests were corrected for multiple comparisons using a Bonferroni correction. Positive correlations are in blue and negative correlations are in red. Shading gives a relative indication of the magnitude of the estimate. ***P ≤ 0.001, **P ≤ 0.01, * P ≤ 0.05. Abbreviations follow Table 1.

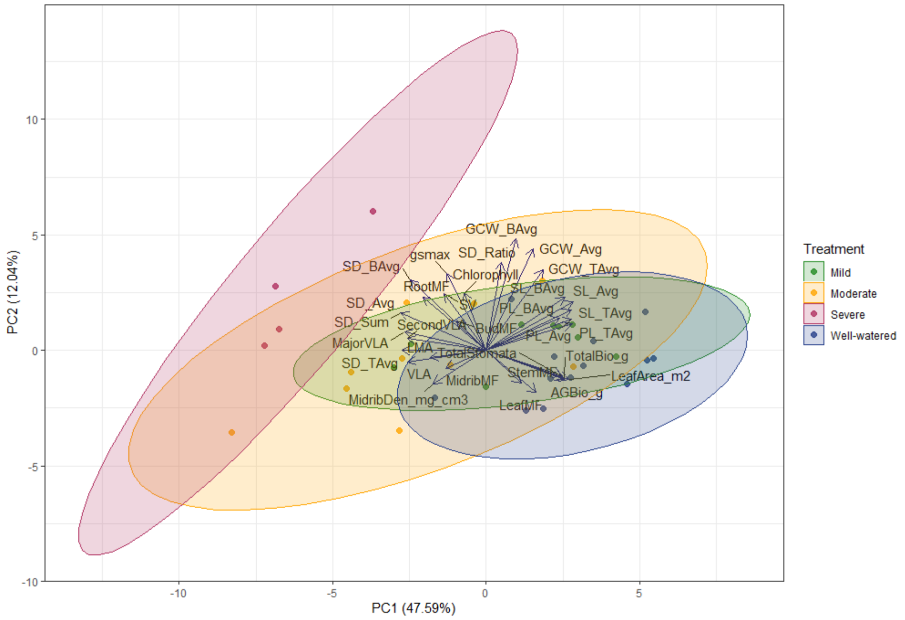


**Figure S3**: Principal component analysis of all traits. Colors indicate treatment as follows: blue = well-watered, green = mild, yellow = moderate, and red = severe. Abbreviations follow Table 1.

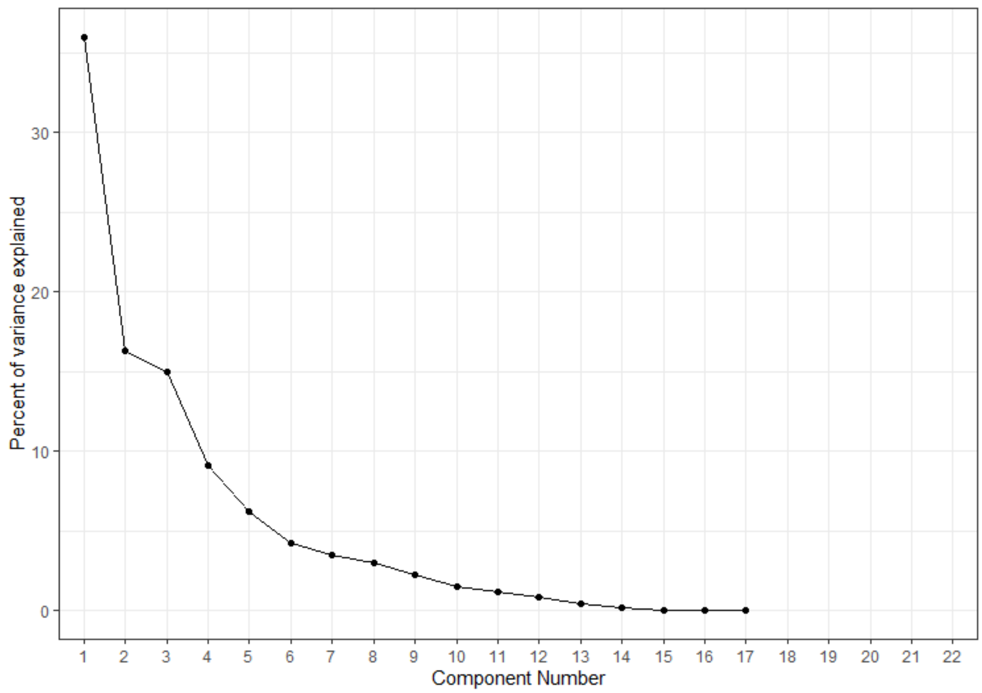


**Figure S4:** Scree plot showing principal components vs. percent variance explained for reduced trait PCA presented in Figure 2.

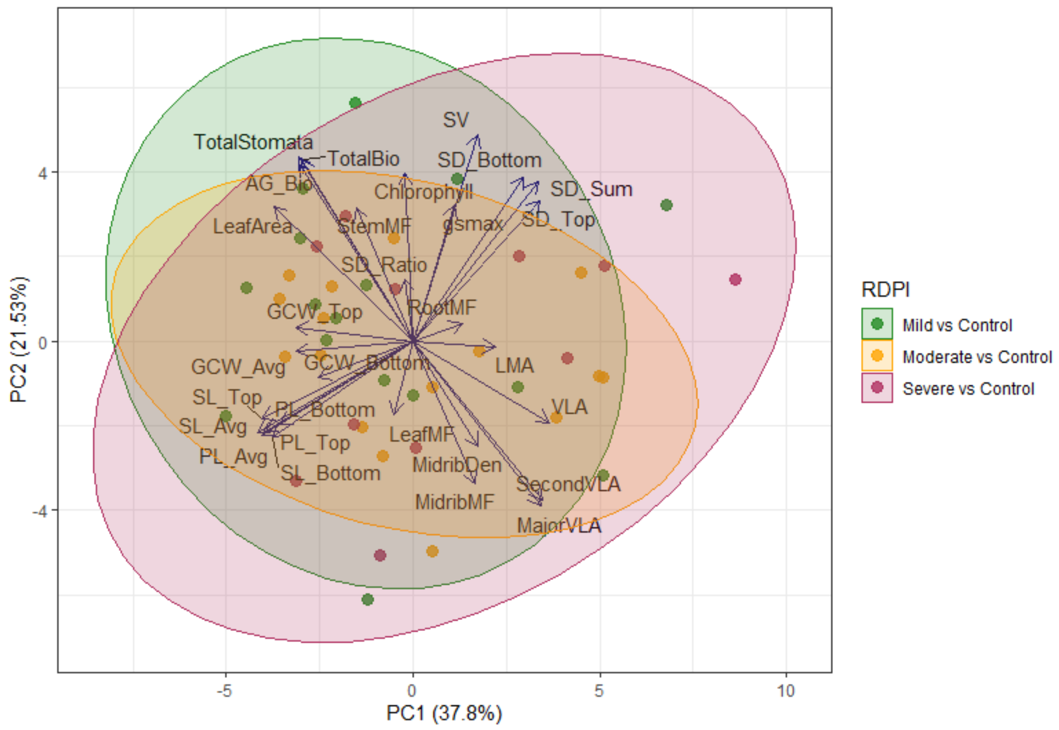

**Figure S5**: Principal component analysis (PCA) of trait plasticity values (estimated as RDPI) for each treatment compared to control for all traits. Abbreviations follow Table 1.

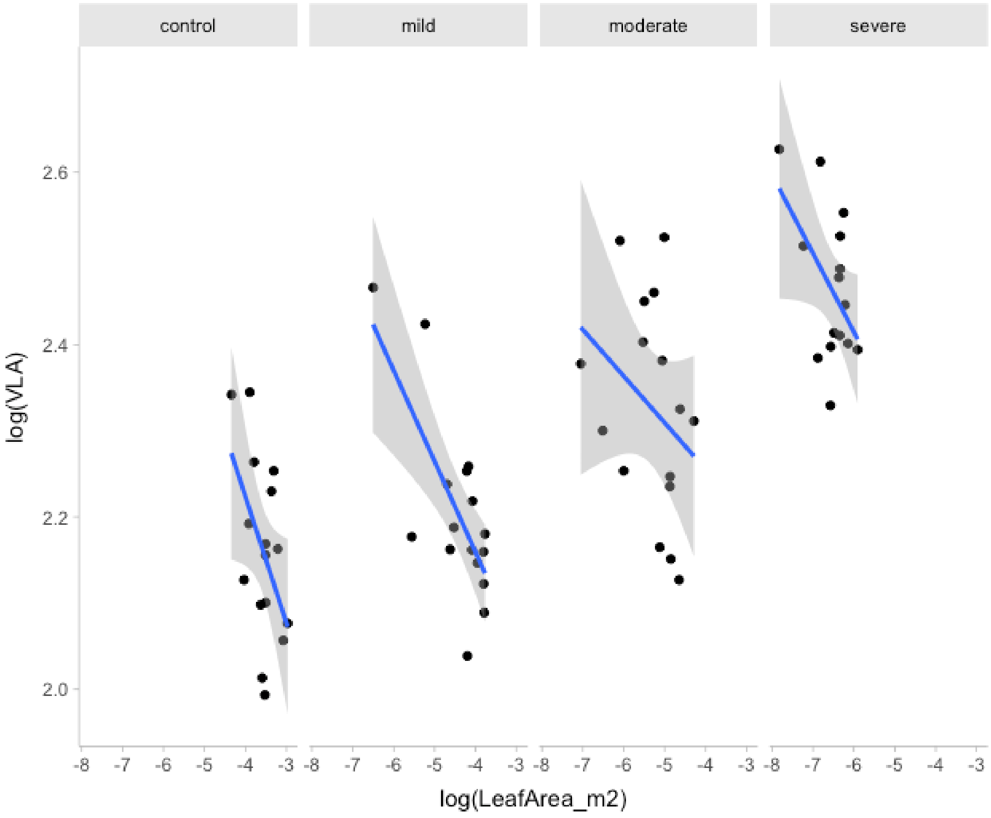


**Figure S6**: Relationship between log transformed VLA and log transformed leaf by treatment. Dots represent individual plants, and the blue line represents the line of best fit. The gray shading is the 95% confidence interval.

**Table S1**: Tukey's HSD test results for all pairwise treatment comparisons. Significance for genotypic effects (***P ≤ 0.001, **P ≤ 0.01, * P ≤ 0.05, P ≤ 0.1, ns = not significant). All traits with significant treatment effects (Table 1) are shown. Traits with no significant treatment effects are excluded from the table.

| Trait | Well watered-Mild | Well watered - moderate | Well watered - Severe | Moderate-Mild | Moderate-Severe | Severe-Mild |
|---|---|---|---|---|---|---|
| SD_Bottom (stomata/mm$^2$) | ns | ns | *** | ns | *** | *** |
| SD_Top (stomata/mm$^2$) | ns | ns | *** | ns | * | *** |
| SD_Sum (stomata/mm$^2$) | ns | ns | *** | ns | *** | *** |
| Stomatal Ratio (bottom/sum) | ns | ns | * | ns | . | ns |
| SL_Bottom (μm) | ns | * | *** | . | ns | *** |
| SL_Top (μm) | ns | * | *** | . | ns | *** |
| SL_Avg (μm) | ns | * | *** | * | ns | *** |
| PL_Bottom (μm) | ns | * | *** | * | ns | *** |
| PL_Top (μm) | ns | . | *** | . | * | *** |
| PL_Avg (μm) | ns | * | *** | * | . | *** |
| Leaf Area (m$^2$) | *** | *** | *** | *** | ** | *** |
| LMA (g/m$^2$) | ns | ns | *** | ns | ** | *** |
| VLA (mm/mm$^2$) | ns | *** | *** | ** | ** | *** |
| 2nd VLA (mm/mm$^2$) | ns | *** | *** | . | ** | *** |
| Major VLA (mm/mm$^2$) | ns | *** | *** | * | ** | *** |
| AG Bio (g) | *** | *** | *** | *** | . | *** |
| Total Bio (g) | *** | *** | *** | *** | . | *** |
| Midrib Density (mg/cm$^3$) | ns | ns | ** | ns | * | ** |
| Midrib MF ($g_{midrib}/g_{leaf}$) | ns | ns | *** | ns | ns | ** |
| $g_{smax}$ (mol/m$^2$s) | ns | ns | . | ns | * | ns |
| Leaf MF ($g_{leaf}/g_{plant}$) | ns | ns | * | ns | ns | ns |
| Stem MF ($g_{stem}/g_{plant}$) | ns | ns | * | ns | ** | * |
| Root MF ($g_{root}/g_{plant}$) | * | . | *** | ns | ** | * |
| TotalStomata (stomata) | *** | *** | *** | *** | . | *** |

**Table S2**: Mean coefficients and credible intervals for trait model shown Figure 5. Abbreviations follow Table 1.

| Trait | Probs | Control_summary | Mild_summary | Moderate_summary | Severe_summary |
|---|---|---|---|---|---|
| VLA | 95_low | -1.04 | -0.98 | -0.24 | -0.08 |
| VLA | 80_low | -0.89 | -0.80 | -0.08 | 0.07 |
| VLA | Mean | -0.61 | -0.48 | 0.18 | 0.38 |
| VLA | 80_high | -0.34 | -0.17 | 0.43 | 0.68 |
| VLA | 95_high | -0.20 | 0.01 | 0.58 | 0.86 |
| SD_Sum | 95_low | -0.17 | -0.73 | -0.04 | -0.19 |
| SD_Sum | 80_low | 0.09 | -0.51 | 0.13 | -0.08 |
| SD_Sum | Mean | 0.63 | -0.09 | 0.43 | 0.15 |
| SD_Sum | 80_high | 1.16 | 0.34 | 0.74 | 0.37 |
| SD_Sum | 95_high | 1.45 | 0.59 | 0.93 | 0.50 |
| SL_Avg | 95_low | -0.41 | -0.71 | -0.07 | -0.20 |
| SL_Avg | 80_low | -0.24 | -0.55 | 0.09 | -0.02 |
| SL_Avg | Mean | 0.07 | -0.27 | 0.37 | 0.31 |
| SL_Avg | 80_high | 0.37 | 0.02 | 0.66 | 0.64 |
| SL_Avg | 95_high | 0.53 | 0.18 | 0.83 | 0.83 |
| logLMA | 95_low | -0.27 | -0.88 | -0.39 | -0.17 |
| logLMA | 80_low | 0.06 | -0.56 | -0.29 | -0.10 |
| logLMA | Mean | 0.66 | 0.01 | -0.08 | 0.03 |
| logLMA | 80_high | 1.29 | 0.54 | 0.12 | 0.16 |
| logLMA | 95_high | 1.60 | 0.88 | 0.23 | 0.24 |
| Biomass | 95_low | 0.39 | -0.95 | -1.36 | -2.33 |
| Biomass | 80_low | 0.81 | -0.56 | -0.93 | -1.86 |
| Biomass | Mean | 1.45 | -0.01 | -0.44 | -1.29 |
| Biomass | 80_high | 2.11 | 0.56 | 0.01 | -0.75 |
| Biomass | 95_high | 2.59 | 1.01 | 0.45 | -0.36 |

**Table S3**: Mean coefficients and credible intervals for RDPI model shown in Figure 6. Abbreviations follow Table 1.

| Probs | VLA_summary | SD_summary | SL_summary | logLMA_summary |
|---|---|---|---|---|
| 95_low | -0.80 | -0.13 | -0.40 | -0.18 |
| 80_low | -0.66 | 0.02 | -0.24 | -0.09 |
| Mean | -0.44 | 0.25 | 0.02 | 0.08 |
| 80_high | -0.22 | 0.48 | 0.28 | 0.25 |
| 95_high | -0.10 | 0.60 | 0.43 | 0.35 |
